# Developmentally Incomplete Barb Rami Increased the Morphological Diversity of Early Feathers

**DOI:** 10.1101/2024.12.07.627359

**Authors:** Yanyun Zhang, Jiawei Tang, Ying Wang, Shuo Wang

**Author notes:** To whom correspondence should be addressed. Shuo Wang., Ying Wang.

## Abstract

Recent studies have significantly advanced our understanding of the evolutionary processes and developmental mechanisms behind feather branching patterns. However, little attention has been given to the tissue differentiation of branches associate with this process. Here, we describe eight feathers preserved in Burmese ambers, characterized by ribbon-like barb rami composed solely of cortex, lacking a medulla. These feathers are categorized into three morphotypes based on the details of their barb rami morphologies. For each morphotype, potential developmental scenarios were proposed through inference of barb development in chickens, suggesting limited tissue differentiation of barb rami in these early feathers. Functional simulations indicate that the configuration of modern barb rami is aerodynamically more stable than those of early feathers, indicating that flexural stiffness may have been a crucial factor driving the evolution of feather branches toward modern configurations. The discovery of developmentally incomplete barb rami in Burmite feathers implies that, while the three-level hierarchical branching pattern of modern feathers had evolved by the Jurassic period, tissue differentiation patterns of feather branches may not have fully stabilized by the Late Cretaceous. This instability likely contributes to the morphological diversity of early feathers, resulting in forms not seen in modern birds.

## 1. Introduction

Extant feathers, with their hierarchical branches of rachis, barbs, and barbules, are among the most complex integumentary structures ever known^1,2^. A typical bipinnate feather consists of a central rachis (feather shaft) extending from the proximal calamus and divides the feather into two vanes^2^. The primary branches of the rachis, called barbs, each have a central ramus that supports two rows of barbules, which are the feather’s secondary branches^1–3^. The morphological diversity of modern feathers arises largely from the branching patterns of the rachis, barbs, and barbules^1,3^, in addition to their coloration.

Except for the calamus and barbules, both the rachis and barbs in modern feathers are characterized by a dense peripheral cortex surrounding a spongy internal medulla^1,4^. This sandwich-like structure grants modern feathers with flexibility and lightness while maintaining the stiffness necessary for flight, body streamlining, and other functions^4,5^. Although studies over the past decades have greatly advanced our understanding of the evolutionary processes and developmental mechanisms behind feather branching patterns^2,3,6–8^, little attention has been given to the evolutionary process of tissue differentiation of feather branches^4,5^. A recent study highlighted the ventrally open rachises documented in Mesozoic feathers, which result from developmental incompleteness of typical cylindrical shafts. This underscores the role of tissue differentiation along the radial axis of the follicle in driving the morphological variations in early feathers^5^. While it remains unclear whether these are pathological structures or simply represent failed attempts of nature, their presence in both non-avialan and avialan theropods, and even in the summer plumage of extant penguins^5^, suggests that incomplete tissue differentiation might have played functional roles in creating rachial variations. These variations may have helped birds adapt to new ecological niches, raising important questions about how the evolution of tissue differentiation of feather branches, in conjunction with branching patterns, has contributed to the morphological diversity of feathers.

Since the rachial ridge forms through the fusion of barb ridges^3^, it has been speculated that ventrally open barb rami may have exist in early feathers, although none had been discovered when ventrally open rachises were first described^5^. Here, we report enigmatic feathers from Burmese ambers, featuring ribbon-like barb rami attached to incomplete rachises. Careful examination suggests that some of these ribbon-like barbs are ventrally open, and barb rami are even split along the midlines in one specimen which create unique mesh-like connections of barbs that have not been documented in extant or fossil birds. Functional simulations reveal that the aerodynamic stability of ramus-split barb is higher than that of barbs with simple ventrally open or plate-like cross-sections in certain condition, though still less stable than the typical sandwich-like configuration of modern feathers. This suggests that these early feathers were likely not adapted for flight or body streamlining. These findings enhance our understanding of how tissue differentiation contributed to the morphological diversity of early feathers, offering insights that may inspire future biomimetic design.

## 2. Materials and Methods

### Institutional Abbreviations

**CNU,** College of Life Sciences, Capital Normal University, Beijing, China; **ECNU,** School of Life Sciences, East China Normal University, Shanghai, China.

### Ancient and extant feathers

All amber-embedded feather specimens described in this paper were donated in 2016 by an anonymous amateur collector to the Biology Museum of East China Normal University (ECNU), Shanghai, China. None of the specimens were associated with Myanmar Economic Corporation, a military-owned conglomerate, ensuring compliance with the moratorium issued by the Society of Vertebrate Paleontology ^9^.

For histological sections, a regenerating 5^th^ primary remex, 4 weeks into regeneration, and an early-stage developing covert contour feather from the breast were extracted from a one-year-old White Leghorn chicken (*Gallus gallus*) that was raised at the animal center of ECNU. The primary remex had reached approximately one fourth of its full growth, and the portion still developing within the follicle was sampled. To label proliferating cells, 1% bromodeoxyuridine (BrdU, Sigma B5002) was injected into the chicken vein 3 hours prior to feather extraction. The procedures for feather collection and paraffin sectioning were conducted in accordance with protocols approved by East China Normal University (Approval Number ARXM2024010).

### Terminology

The terms used to describe feather structures and orientations follow Lucas and Stettenheim (1972)^1^, while the descriptions of the developing follicle adhere to Prum (1999)^3^. Specifically, “proximal” and “distal” refer to structures closer to or farther from the calamus, respectively; while “central” and “peripheral” refer to structures near or away from the follicle center, respectively. Throughout the paper, we employed terms like “sandwich-like,” “ribbon-like,” “plate-like,” and “beam-like” to describe different configurations of feather barbs. “Sandwich-like” refers to barb rami typical of modern feathers, where an internal spongy medulla is surrounded by a cortex layer when viewed in cross-section. “Ribbon-like” describes barb rami with only a dorsal cortex, showing some differentiation of dorsal cortex but lacking a distinct medulla and ventral cortex. “Plate-like” is used for barb rami that consist solely of an expanded cortex, without distinct medulla; while “beam-like” indicates the most primitive configuration, featuring minimal differentiation of the dorsal cortex and no other tissue specialization.

### Histological sections and staining

The growing portion of the regenerating chicken primary remex was embedded in paraffin and sectioned at 5 mm intervals, starting from the immature proximal end to establish a growth series. After 4 hours of fixation in 4% paraformaldehyde, samples were dehydrated through a graded ethanol series ranging from 30% to 100%. Paraffin sections with a thickness of 7μm were cut perpendicular to the feather’s long axis. Hematoxylin and Eosin (H&E) staining followed established protocols by Fischer et al. (2008)^10^, and BrdU assays were performed according to Lee et al. (2001)^11^, using an AEC Immunohistochemistry Color Development Kit (Sangon Biotech, E670031) instead of the chromogenic method^11^. Masson staining was conducted at developmental stage I primary remex following Suvik (2012) to complement the H&E stanning^12^. Digital images of histological slides were captured using a Nikon Digital Sight 10 Camera System attached to a Nikon Ni-E transmitted microscope.

### Micro CT scanning

Computed tomography (CT) scans of ECNU A32 were performed using a beam energy of 18kV at a resolution of 4.0um per pixel using the Phoenix v-tome-x industrial CT scanner at the Molecular Imaging Center, University of Southern California. ECNU A143-1 was scanned under 12 kV at a resolution of 3.9um per pixel using the same scanner at Shanghai Yinghua Inspection & Testing. Segmentation was carried out utilizing Mimics 15 and Amira software at the University of Southern California.

### Functional Simulation

The flexural stiffness of feathers with different cross-sectional shapes of barb rami were estimated using ANSYS FLUENT 12.0 software^13^. 3D models of ECNU A32 and an uncategorized contour feather of similar size from a one-year-old White Leghorn chicken were compared to assess the aerodynamic performance (Fig. S1A, B). Due to limited computing power, simplified models were used for aerodynamic analyses of feathers with various barb ramus cross-sections (Fig. S2). Barbules are omitted from the simplified models because they are not the focus of the present study. A ribbon-like branch with crescent-shaped cross-section simulate the ventrally open barb rami of morphotype I feathers (Fig. S2A, model I); a ribbon-like branch with paired half-crescents cross-section simulate morphotype II feather barbs, where barb ramus are made-up by halves of two adjacent split barb branches (Fig. S2B, model II); a branch with a 10:1 flat elliptical solid cross-sections simulate the plate-like barb ramus of morphotype III feathers (Fig. S2C, model III); a branch with a perfectly round and hollow cross-sections simulate idealized, non-realistic barb ramus (Fig. S2,D, model IV), and a branch with a 5:1 elliptical hollow cross-sections simulate the barb ramus of modern flight feathers (Fig. S2E, model V). Structural parameters of each model are given in Fig. S3.

To investigate whether the aerodynamic performance of a solid cross-section differs from that of hollow cross-section with the same parameters, simulations also included branches with a perfectly round solid cross-section (Fig. S2F, model VI) and a 5:1 elliptical solid cross-section (Fig. S2G, model VII) were also included in the simulation. For simplicity, the rachis in each model was simplified to a cylindrical shape and completely fixed, with only the barbs free to move. Airflow was assumed to be evenly applied. Since the barbs were not interlocked, no complete vane was simulated in the analyses.

### Fluid-Structure Interaction (FSI) Analysis

Fluid-structure interaction (FSI) analysis examines how deformable solid structures behave under fluid flow forces and how structural displacement affect fluid flow^14^. Due to the minimal deformation of feather under airflow, its impact on the flow field distribution was considered negligible, allowing for a one-way FSI approach^15^. The FSI analysis include: 1) modeling the external flow field surround the feather; 2) transferring the results of the flow field analysis to the structural field; and 3) performing one-way coupled analysis. Since the barbs and rachis are composed of feather keratin, material properties were set based on real feather keratins, with a density of 1.15g/cm^3 16^, a Young’s modulus (a parameter characterizing the material stiffness against the external force) of 2.5× 10^9^ N/M^2 17,18^, and a Poisson’s ratio of 0.45^19^. Each model feather was set to a length of 60 mm, with 10mm long barbs forming a 45° angle with the rachis (Fig. S2 A). For all models, the cylindrical rachis had a diameter of 2 mm, and the barb rami had a diameter of 1 mm, with a cortex thickness of 0.05 mm for hollow rami (Fig. S3). The ventrally open barb rami of morphotype I feathers corresponded to half of a hollow ramus (Fig. S3①), while each of the paired ramus branches of morphotype II feather corresponded to a quarter of the hollow ramus (Fig. S3②).

Due to the significant differences in elasticity between the cortex and medulla, the flexural stiffness of feather barbs is primarily determined by the geometry of the cortex^20–22^. Simulations subjected each morphotype to airflow either perpendicular to the feather vanes or parallel to the feather rachis from the calamus, exploring aerodynamic performances under lateral and headwind conditions, respectively. Since the primary function of covert contour feathers is to streamline the body surface by allowing airflow to pass smoothly along the feather from the calamus, we believe the headwind condition closely replicate the aerodynamic performance of contour feathers in their natural state. While absolute values may not reflect real-world conditions, the relative flexibility (displacement from the static state) and the static stress responses to standardized forces offer insight into the aerodynamic performance of each feather type.

## 2. Results

### Morphotypes

Eight feathers preserved in seven pieces of amber are included in this study. While rachial morphology varies both among feathers and along their lengths, all specimens show evidence of a ventrally open rachis, although the extent varies. For example, in ECNU A143-1 (Fig. 1A), the ventrally open region extends throughout the entire rachis similar to that of CNU A0005^5^, whereas in ECNU A147, it is restricted to the proximal half (Fig. 1J). Since this study focuses primarily on the barbs rather than the rachis and barbules, the feathers are categorized into three morphotypes based on the morphology of the barb rami:

**Figure 1.**
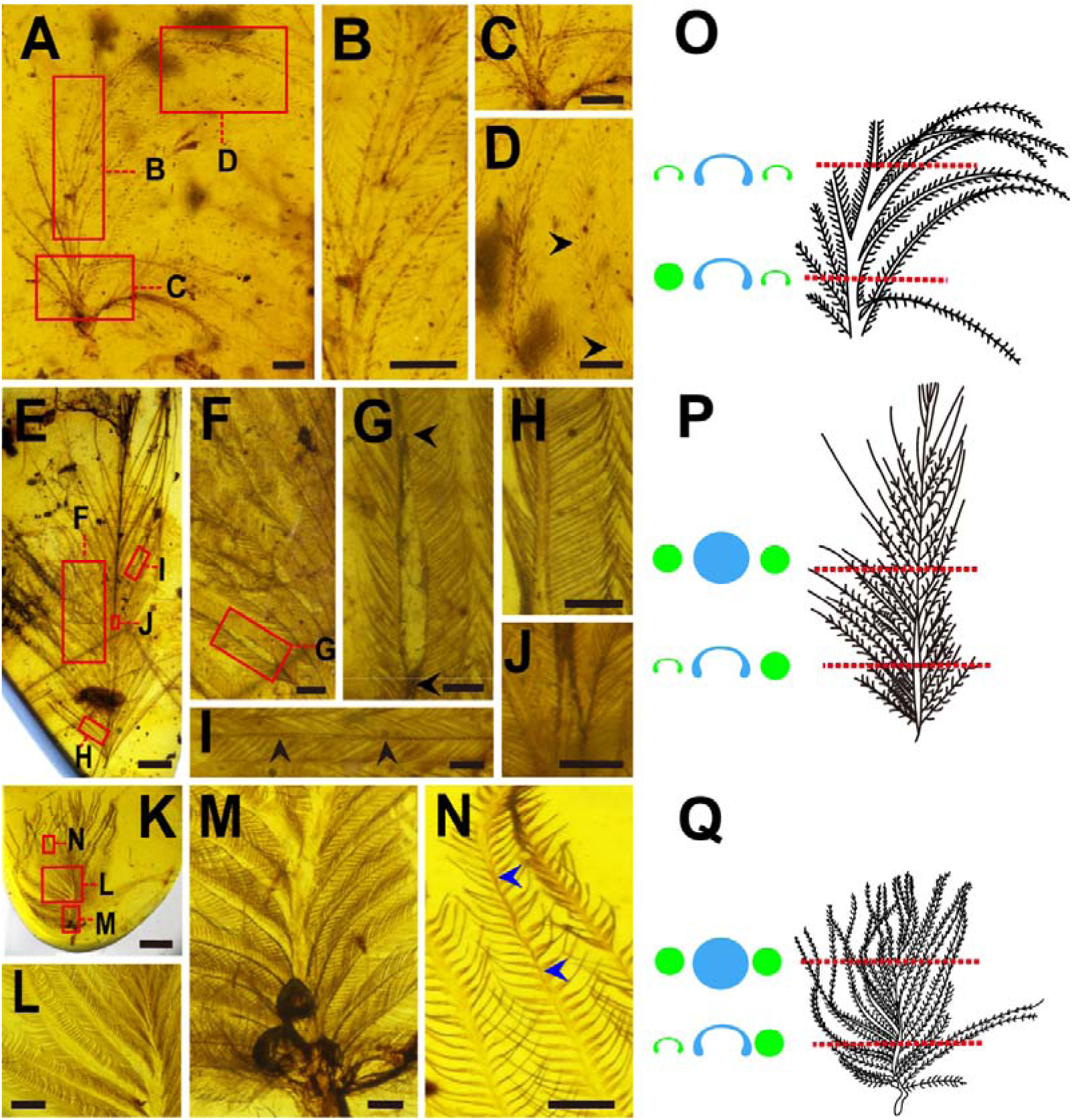
Morphotype I feathers showing ventrally open barb rami. **Left (A-L):** Photographs of isolated feathers showing ventrally open barb rami: (A-D) images of ECNU A143-1, (E-J) images of ECNU A147, (K-N) images of ECNU A148; red boxes in (A) indicate the areas shown in close-up images (B-D); red boxes in (E) and (F) indicate the areas shown in close-up images (F, H, I, J) and (G), respectively; red boxes in (K) indicate the areas shown in close-up images (L-N). Paired black arrowheads in (D) mark the position where barb ramus is ventrally opened, whereas paired blue arrowheads in (N) highlight where solid barb ramus is present. **Right (O-Q),** Schematic drawings illustrating general morphologies (right) and cross-sections of rachis and barb rami (left) at various positions along the corresponding feather. Red dashed lines indicate where cross-sections were taken, blue silhouettes represent solid (circle) or ventrally open (crescent-shaped) rachial cross-sections, while green silhouettes indicate solid (circle) or ventrally open (crescent-shaped) barb rami cross-sections on either side of the rachis (not to scale). Scale bar: 0.5mm for (A), (E), (F), and (M); 0.2mm for (B-D), (G-J), and (N-L); and 1.5mm for (K). (O-Q) are not to scale.

**Morphotype I** (Figs. 1-2) is represented by four amber specimens: ECNU A49, A143, A147, and A148. ECNU A147 and A148 each contain a nearly complete isolated feather with symmetrically distributed barbs and barbules. ECNU A143 includes two isolated feathers and are labeled as ECNU A143-1 (Fig. 1A-D) and A143-2 (Fig. 2A-D), respectively. ECNU A49 includes only a single barb (Fig. 2E-G), thus little is known about the rest of the feather. All morphotype I feathers are characterized by proximodistally broad, ribbon-like barb rami with barbules along their proximal and distal edges. CT images reveal that these ribbon-like barb rami consist solely of a thin dorsal cortex that is crescent-shaped in cross section, without presenting a medulla or a ventral cortex (Fig. S4). This absence results in ventrally open rami that have not been observed in modern birds. However, the ribbon-like rami lack a midline ridge and lateral flanges seen in the previously reported ventrally open rachises^5^, suggesting they are mechanically more flexible relative to the developmentally incomplete ventrally open rachises (see ***Aerodynamic performance of different barb rami*** for more information).

**Figure 2.**
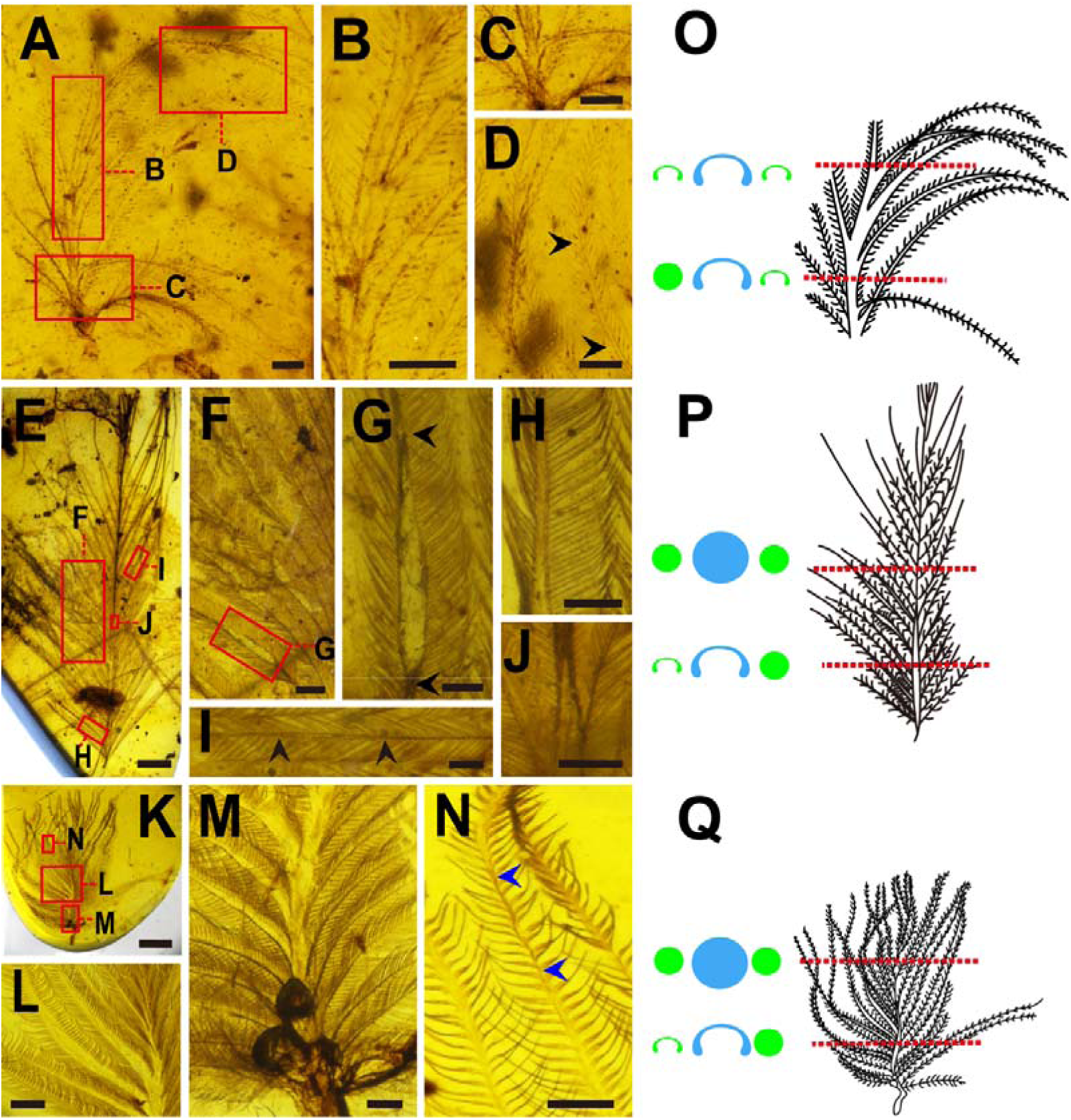
Morphotype I feathers showing ventrally open barb rami. **Left (A-G):** Photographs of isolated feathers showing ventrally open barb rami: (A-D) images of ECNU A143-2, (E-F) images of ECNU A49. Red boxes in (A) indicate the areas shown in close-up images (B-D); while red boxes in (E) highlight areas shown in close-up images (F-G). Paired black arrowheads in (C) mark the position of the ventrally open barb ramus. **Right (H-I),** Schematic drawings illustrating general morphologies (right) and cross-sections of rachis and barb rami (left) at various positions along the corresponding feather (H) or barb (I). Red dashed lines denote the locations from where cross-sections were taken, blue silhouettes represent ventrally open (crescent-shaped) rachial cross-sections, and green silhouettes indicate solid (circle) or ventrally open (crescent-shaped) barb rami cross-sections (not to scale). Scale bar: 0.5mm for (A) and (E); 0.2mm for (B), (D) and (G), 0.1mm for (C) and (F). (H-I) are not to scale.

Ribbon-like barb rami are found in specific regions of individual barbs or across different barbs along a feather shaft. For instance, in ECNU A148 (Fig. 1L, blue arrowheads), ribbon-like regions alternate with solid beam sections, while in A143-1 (Fig. 1A, O), most barb rami are ribbon-like proximally and transition into solid beams distally. This variation reflects the differential expansion of the dorsal cortex during the development of a single barb. Therefore, **morphotype I** feathers are characterized by ribbon-like, ventrally open barb rami, with barbules present along their proximal and distal edges.

**Morphotype II** (Fig. 3) is represented by a single isolated feather (ECNU A32), which features nearly symmetrically distributed barbs and a relatively thick rachis (Fig. 3A-I). CT images show that the rachis has a ventral cleft with a V-shaped cross-section, rather than circular or crescent-shaped one (Fig. 3B, K-M). The barbs along the proximal and distal quarters of the rachis are beam-like, but those in the middle form forks soon after branching off from the rachis (Fig. 3J and J’). The fork branches are ribbon-like, each corresponding to half of the crescent-shaped dorsal cortex of a barb ramus. Distally, the branches from adjacent split barbs fuse to form a solid beam, resulting in a distinctive mesh-like structure (Fig. 3Q). Notably, barbules are absent along the edges of the barb rami but present on the fork branches, where they are oriented toward the center of the fork (Fig. 3J and J’). This suggests that the fork branches represent halves of adjacent barb rami that split along the midline (Fig. 3N-Q), a configuration not observed in any other fossil or modern feathers. In cross-section view, the distal fork branch consistently lies ventrally to the proximal branch where they merge into the rachis (Fig. 3L-M), giving the impression that barbs twist immediately after branching from the feather shaft (Fig. 3J-K). Thus, **morphotype II** feathers are distinguished by the presence of forked barb rami with barbules along the fork branches, oriented toward the center of the fork.

**Figure 3.**
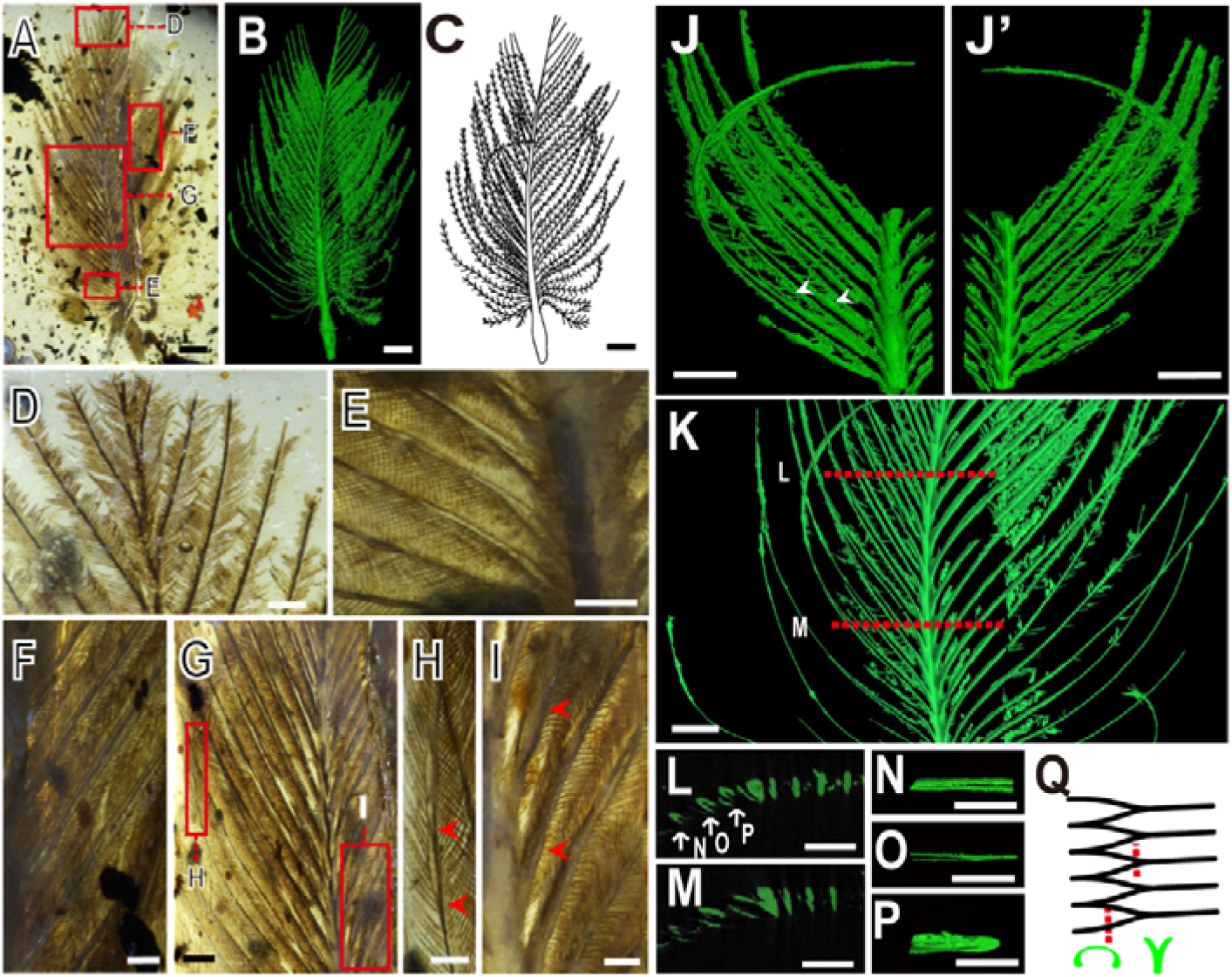
Morphotype II feathers showing the split ventrally open barb rami. Photograph (A), 3D segmentation (B) and schematic drawing (C) of the isolated feather ECNU A32. Red boxes in (A) highlight areas shown in close-up images (D-H), while red boxes in (F) indicate areas shown in close-up images (G) and (I). Paired black arrowheads in (G) and (I) indicate positions where the ventrally open barb rami are present. Representative sections (J-P) from the 3D segmentation of ECNU A32 highlight the split barb rami (red arrowheads) in dorsal (J) and ventral (J’) views. Red lines in (K) mark the positions where the cross-sections shown in (L) and (M) were taken. (N-P) each highlights a barb fork, which connect to form a unique mesh-like structure (Q, not to scale), with red lines marking the positions of cross-sections of the barb rami shown as green silhouettes. Scale bar: 1.25mm for (A-C) and (J-P); 0.2mm for (D-I’).

**Morphotype III** (Fig. 4) is represented by two amber specimens, ECNU A101 (Fig. 4A-E) and A149 (Fig. 4F-I), each containing a single feather with apparently symmetrical distribution of barbs and barbules. Unlike other feather morphotypes, the barb rami of Morphotype III feathers are compressed proximodistally and broad dorsoventrally (Fig. 4D, I). There is no medulla present, and barbules are distributed along both the distal and proximal surfaces of the barb rami. Thus, **morphotype III** feathers are distinguished by the absence of a medulla and the presence of proximodistally compressed barb rami with barbules attached to both surfaces, setting them apart from morphotype I feathers.

**Figure 4.**
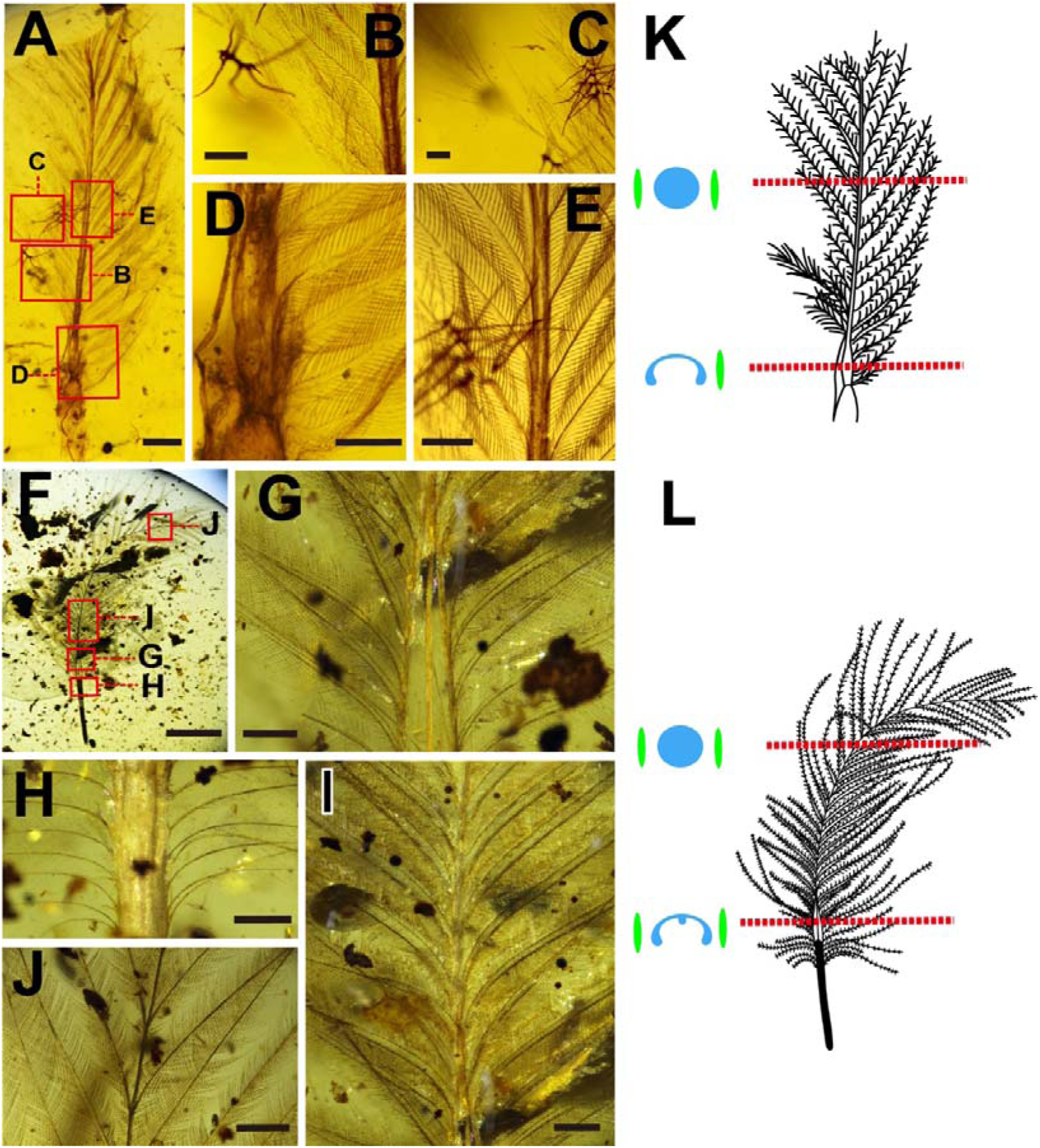
Morphotype III feathers showing plate-like barb rami without medulla. **Left (A-I):** Photographs of isolated feathers showing plate-like barb rami: (A-E) images of ECNU A101, (F-I) images of ECNU A149. Red boxes in (A) indicate the areas shown in close-up images (B-E); while red boxes in (F) highlight areas shown in close-up images (G-J). **Right (K-L),** Schematic drawings illustrating general morphologies (right) and cross-sections of rachis and barb rami (left) at various positions along the corresponding feather. Red dashed lines mark the locations where cross-sections were taken, blue silhouettes represent solid (circle) or ventrally open (crescent-shaped) rachial cross-sections, and green silhouettes indicate plate-like barb rami cross-sections on either side of the rachis (not to scale). Scale bar: 0.5mm for (A) and (G-I); 5mm for (F); 0.2mm for (B-E). (K-L) not to scale.

### Aerodynamic performance of different barb rami

Our simulation results show that, among all tests, feather barbs with a perfectly round cross-section exhibit the highest strength, while those with elliptical cross-sections demonstrate relatively lower strength (Figs. 5-6). This suggests that the flexural stiffness of feather barbs is primarily determined by cross-sectional geometry, rather than the material properties of feather keratins^4,18,23^. In both sets of tests, barb rami with solid cross-sections consistently show less deformation compared to those with hollow cross-sections (Figs. 5-6), indicating that a solid structure, given the same shape and diameter, significantly enhances mechanical strength.

**Figure 5.**
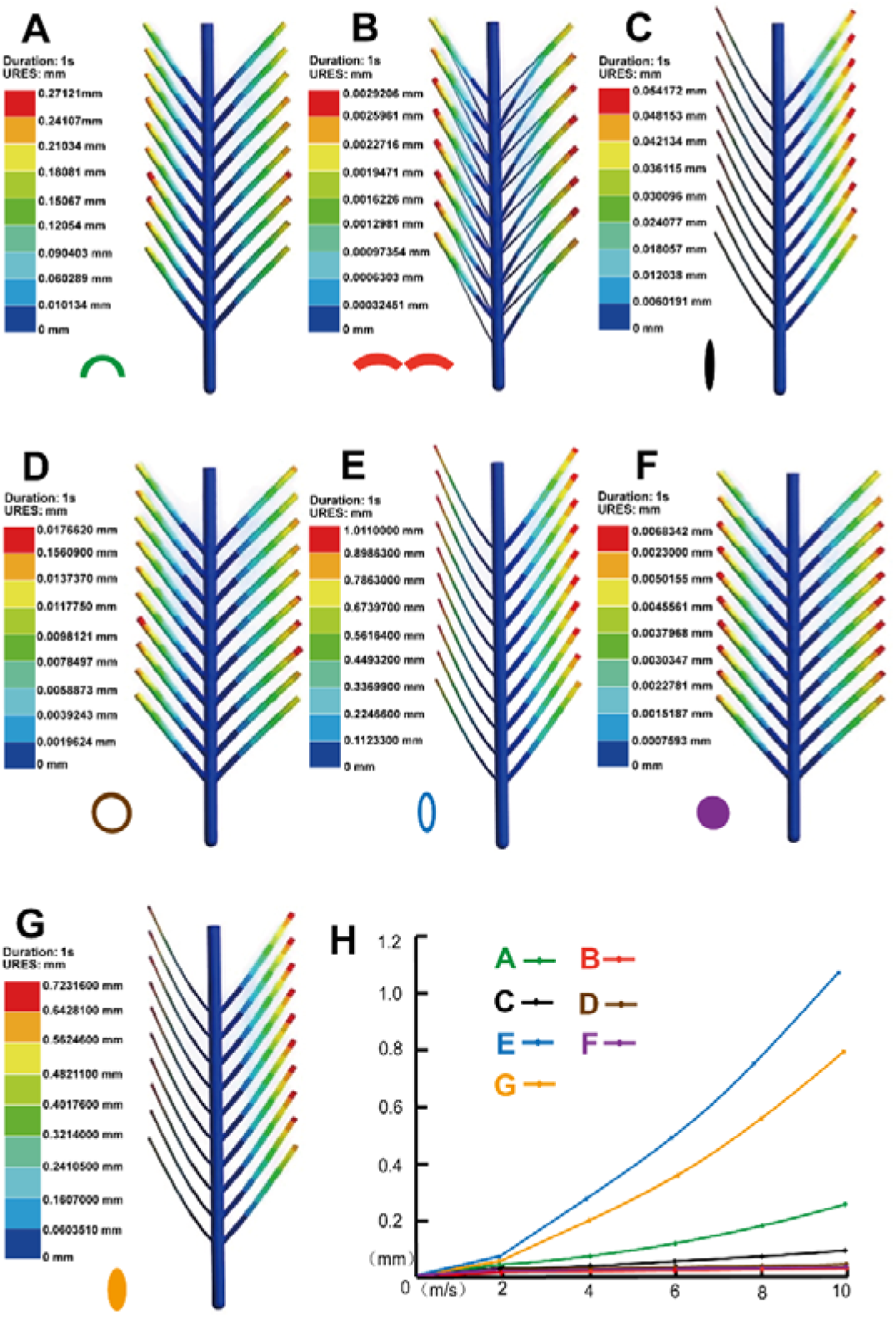
Simulations illustrating the maximum displacements of feather barbs with various cross-sectional shapes under airflow perpendicular to the feather vanes. The models simulate: (A) morphotype I feather barb rami with crescent-shaped cross-sections; (B) morphotype II feather barb rami with paired half-crescents cross-sections; (C) morphotype III feather barb rami with 10:1 flat elliptical solid cross-sections; (D) idealized, non-realistic tubular barb rami with perfectly round and hollow cross-sections; (E) modern flight feather with 5:1 elliptical hollow barb rami cross-sections; (F) idealized, non-realistic tubular barb rami with perfectly round and solid cross-sections; and (G) idealized, non-realistic barb rami with 5:1 elliptical solid cross-sections. Resultant displacement (URES) is expressed in millimeters, with the symbol next to each model indicating the shape of barb ramus cross-sections. A comparison of aerodynamic performance of each simulated model is shown in (H), with the *x*-axis representing airflow speed and the *y*-axis representing displacement. The results demonstrate when airflow is perpendicular to the feather vanes, barbs with paired half-crescent cross-section exhibit the greatest strength (B, H), whereas barbs with a 10:1 flat elliptical solid cross-section exhibit the least strength (C, H). This highlights the significant impact of cross-section shape on barb flexibility and deformation under perpendicular airflow.

**Figure 6.**
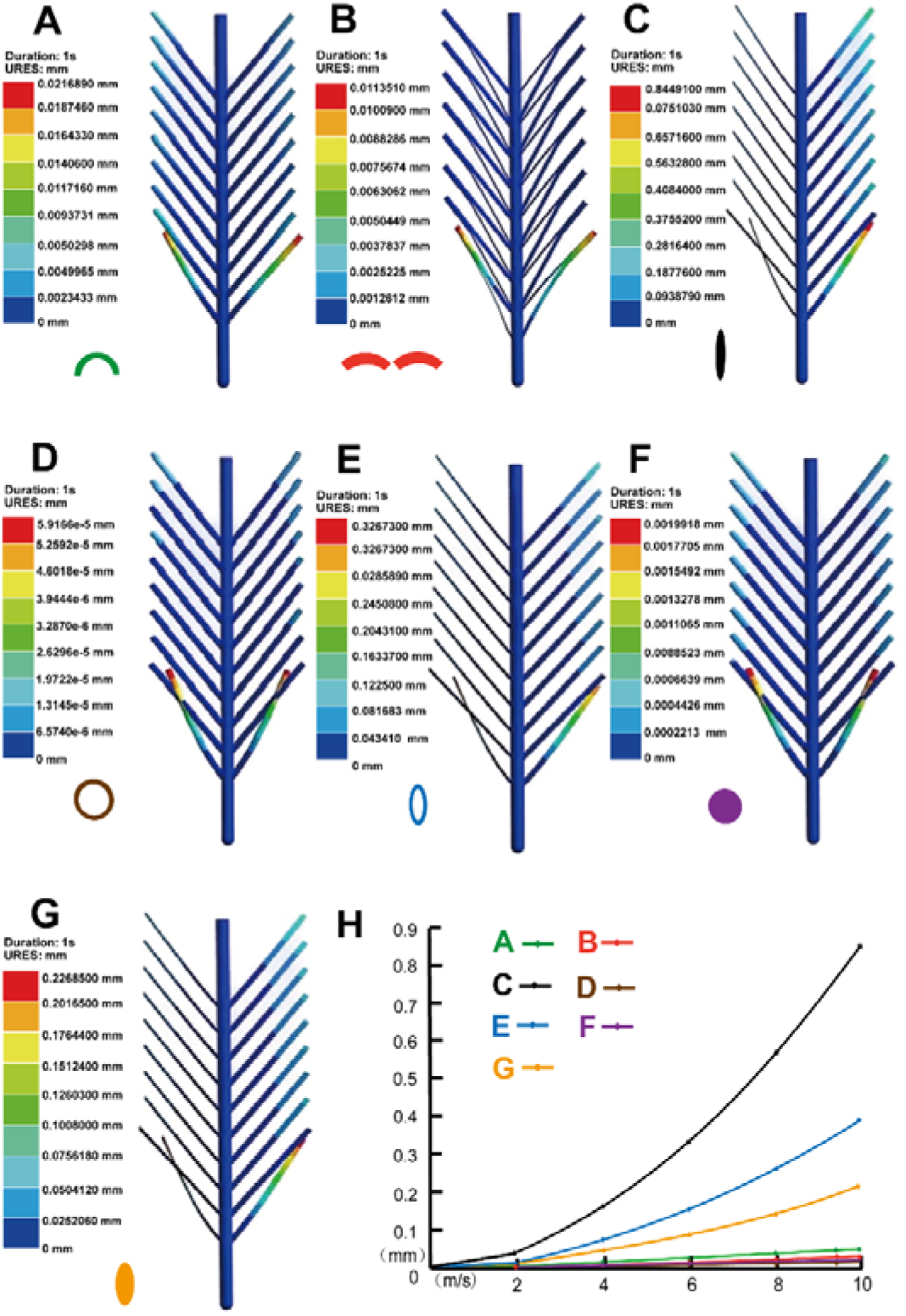
Simulations illustrating the maximum displacements of feather barbs with various cross-sectional shapes under airflow passing along feather vanes from the calamus. The models simulate: (A) morphotype I feather barb rami with crescent-shaped cross-sections; (B) morphotype II feather barb rami with paired half-crescents cross-sections; (C) morphotype III feather barb rami with 10:1 flat elliptical cross-sections; (D) idealized, non-realistic tubular barb rami with perfectly round and hollow cross-sections; (E) modern flight feather with 5:1 elliptical hollow barb rami cross-sections; (F) idealized, non-realistic tubular barb rami with perfectly round and solid cross-sections; and (G) idealized, non-realistic barb rami with 5:1 elliptical solid cross-sections. Resultant displacement (URES) is expressed in millimeters, with the symbol next to each model indicating the shape of barb ramus cross-sections. A comparison of aerodynamic performance of each simulated model is shown in (H), with the *x*-axis representing airflow speed and the *y*-axis representing displacement. The results demonstrate that when airflow passes along feather vanes from the calamus, barbs with a hollow cross-section are the most stable configuration (D, H), whereas barbs with a 5:1 flat elliptical hollow cross-section exhibit the least strength (E, H). This highlights the significant impact of cross-section shape on barb flexibility and deformation under parallel airflow.

Specifically, feather barbs with a perfectly round and hollow cross-sections represent the most stable configurations when airflow passes along the feather vanes from the calamus (Fig. 6D). However, when airflow is perpendicular to the feather vanes, forked barbs with paired half-crescent cross-section exhibit greater strength (Figs. 5B, 7A). This advantage arises not from increased resistance of the paired half-crescent shape itself, but because these fork branches connect to form a mesh-like structure, providing better stability under perpendicular airflow compared to other cross-sectional shapes. When subjected to lateral airflow, barb rami with a 1:10 flat elliptical cross-section experience less displacement than those with a 1:5 elliptical cross-section (Fig. 5H), due to the smaller windward area of the former. However, when airflow is directed from the calamus, barbs with a 1:10 flat elliptical cross-section exhibit the greatest displacement (Fig. 6H), as their larger windward area causes more deflection compared to the 1:5 elliptical cross-section.

**Figure 7.**
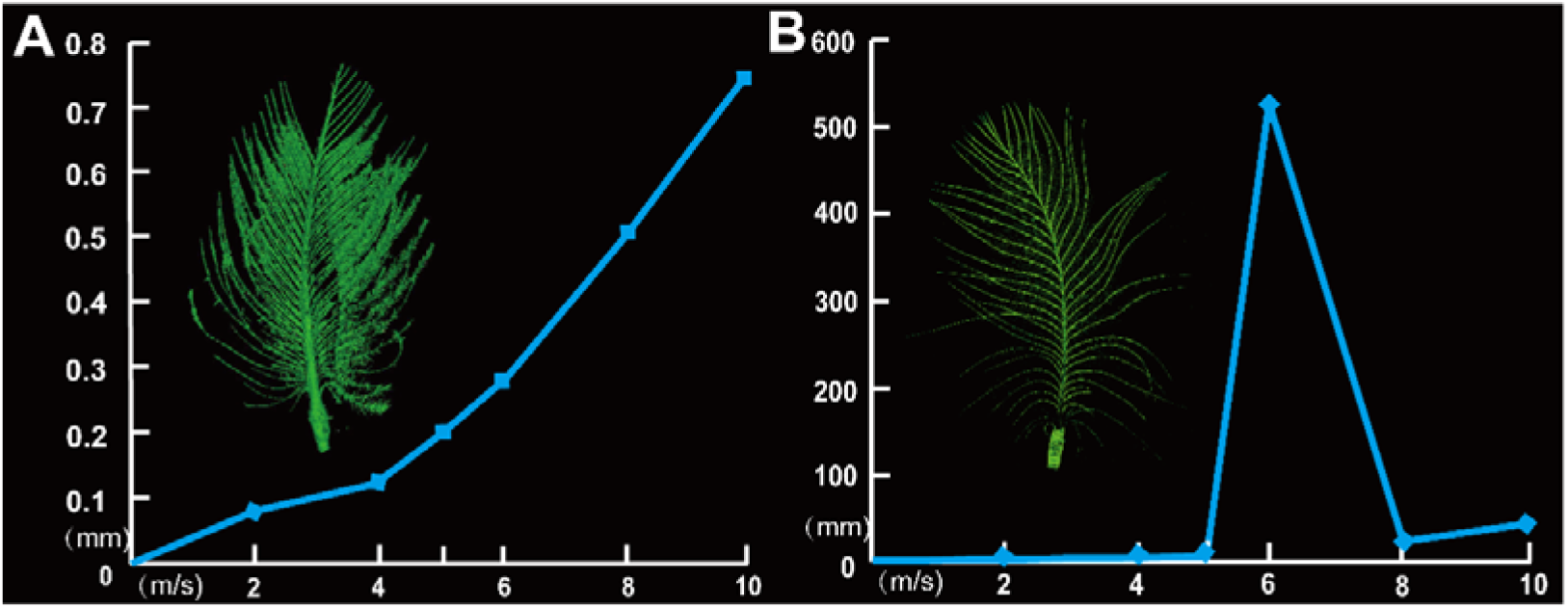
Aerodynamic performance of ECNU A32 (A) and an extant contour feather of similar size (B) under increased airflow, showing the different in displacement cause by structural variations. The *x*-axis represents airflow speed, and the *y*-axis represents displacement. The results show that the basic structure of the extant contour feather is completely destroyed when the airflow speed reaches 6 m/s, leading to an unrealistic displacement of 550 mm. In contrast, no apparent structural destrusction is observed in the ECNU A32 under a maximum airflow speed of 10m/s, demonstrating that the amber-embedded feather with forked barbs exhibits greater mechanism strength.

These findings indicate that barb rami with a 1:5 elliptical cross-section, designed to simulate modern feather barbs, are among the most unstable structures (Figs. 5H and 6H). Nonetheless, modern feather barbs overcome this structural limitation by interconnecting into feather vanes through hooklets, forming an interlocking structure that is functionally more advanced than the mesh-like connections observed in ECNU A32. This interlocking enhances mechanical strength, allowing modern feathers to withstand airflow forces while minimizing self-weight.

### The making of a typical feather barb

A barb, the primary branch of a feather, consists of a ramus (main shaft) lined with rows of barbules on either side^2^. Barbs from different feathers or various positions along a single feather (e.g., covert contour) can exhibit diverse cross-sectional shapes^1^, which may also vary along a single barb^24^. Although most known extant feather barbs display a proximodistally compressed, sandwich-like cross-section with an outer cortex and an internal medulla^4^, not all barbs or all sections of a single barb contain a medulla, depending on the species and feather type.

The development of typical feather barbs follows a highly organized process involving elongation of barb filaments and cellular differentiation along these filaments^2,25^. Initially, a few barb ridges form on either side of the rachial ridge, followed by new ridges arising aside all the way down to the new barb locus, which is located opposite, though not strictly, to the rachial ridge (Fig. 8B, Stage I). Unlike the stationary rachial ridge, the barb ridges move both axially (parallel to the long axis of the follicle) and tangentially (perpendicular to the long axis of the follicle) during growth as existing ridges are carried away^1,3^.

**Figure 8.**
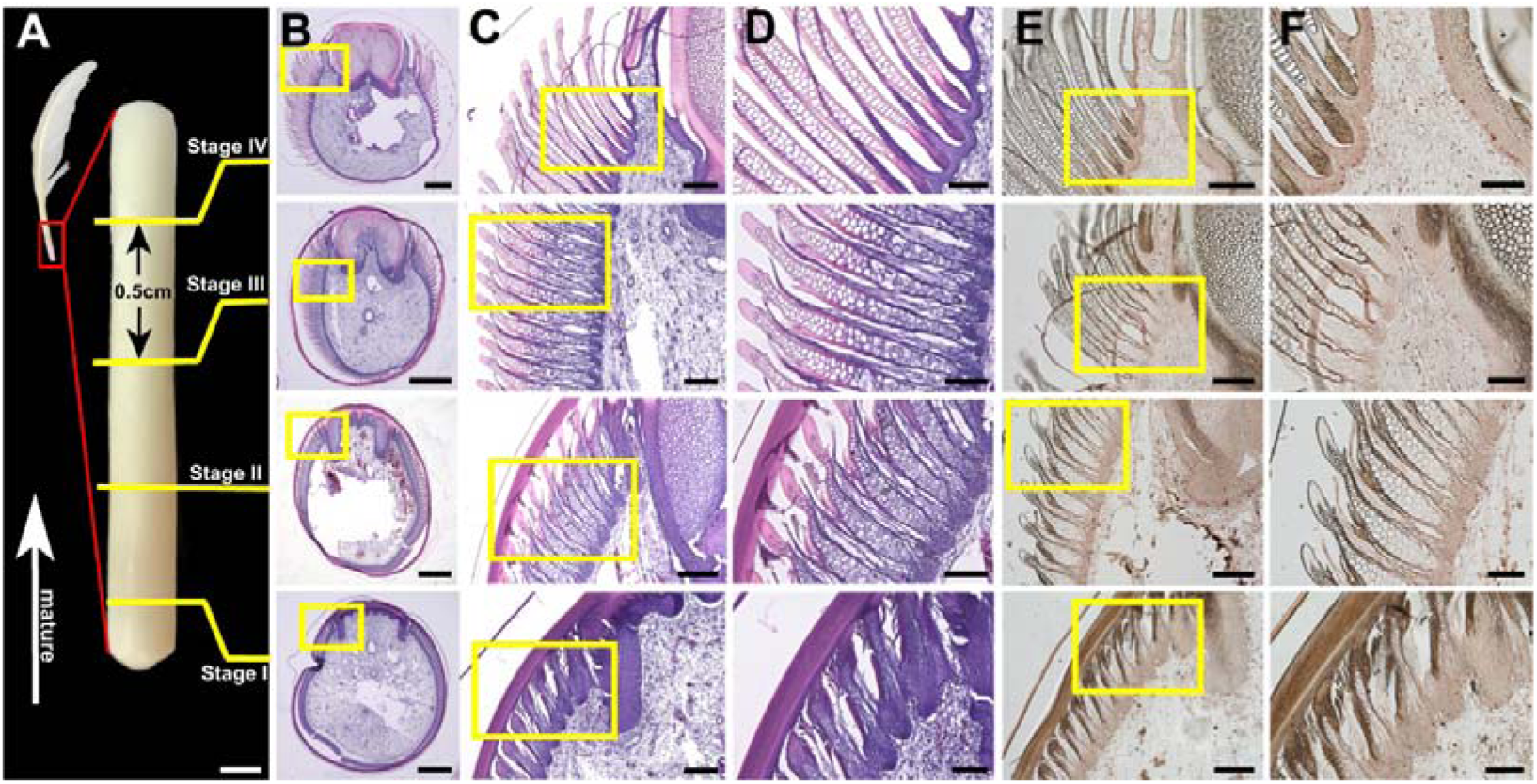
Tissue differentiation (H&E) and cell proliferation (BrdU) of a regenerating 5^th^ primary remex from a one-year old chicken. **The p**hoto of the regenerating remex is shown in (A), with the red box indicating an enlarged image of the sampled developing portion within the follicle, and the yellow lines mark positions of slices shown in (B-F). (B-D) H&E staining, illustrating barb morphologies at different levels of the regenerating remex; and (E-F) BrdU staining, highlighting cell proliferation at each level of a regenerating remex. Yellow boxes mark the areas shown in the corresponding close-up images. Scale bar: 2mm for (A); 1mm for (B); 0.2mm for (C) and (E); 0.1mm for (D) and (F)

Axially, in a cross-section truly perpendicular to the follicle’s long axis, all barb ridges appear at roughly the same developmental stage since they grow at an equal rate (Fig. 8B). This uniformity enables spatiotemporal changes in the developing feather to be characterized^1^. Meanwhile, tangential movement causes the morphology of the rami and barbules to vary along a single barb due to the concentration gradients of several morphogens across the pulp periphery^4^ (Fig. 8B, Stages II and III). The combination of the axial and tangential movements result in a helical displacement of barb ridges toward the rachial ridge^1^ (as seen in serial cross-sections in Fig. 8), ultimately integrating the proximal ends of the barb ridges into the rachial ridge (Fig. 8B, Stage IV).

Histological assays, including H&E, Masson, and BrdU staining of serial cross-sections from developing chicken remex follicles, reveal that at the cellular level, basilar cells in the central part of the epidermal collar are initially surrounded by a thicker layer of intermediate cells (Fig. S5 A-C). These intermediate cells soon reorganize into barb ridges along the radii of the follicle, separated from one another by shallow furrows intercepted by pulp bulges (Fig. S5 D-I). Shortly after their formation, barb ridges begin to differentiate into the various components of barbs. Within each barb ridge, a single layer of basilar cells appears on either side of the intermediate cells, creating the marginal plates (Fig. S5 D-I). The remaining intermediate cells aggregate into an axial plate in the middle, flanked by barbule plates on each side, and terminates centrally at the ramus (Fig. S5 F, I).

Cell proliferation in the barb ridges is not localized in clusters, as BrdU-positive cells are scattered throughout the epidermal basal layer, appearing not only at the tips of barb ridges but also within the furrows (Fig. 8E-F. Stage I). As the barb ridges move tangentially toward the rachial ridge, the barbule plates rapidly differentiate and are the first to undergo keratinization during feather morphogenesis (Fig. 8, Stage I). Apoptosis of the axial plate releases mature barbules, while apoptosis of the marginal plates separates the barb ridges from one another. Keratinization soon extends from the barbule plates to the peripheral walls (equivalent to the dorsal cortex of mature barb rami) of the developing barb rami (Fig. 8, Stage II). At this stage, a high-proliferation zone marked by more BrdU-positive cells appears along the epidermal basal layer (Fig. 8E-F, Stage II), indicating that the barb ridges are growing toward the follicle center, similar to the rachial ridge^5^.

As the expanding medullae push the barb rami all the way toward the peripheral feather sheath, keratinization spreads along the lateral cortex to the central walls (equivalent to the ventral cortex of mature barb rami) of the developing barb rami (Fig. 8, Stage III). At this point, cell proliferation decreases, and the medullae become fully keratinized once the peripheral walls of the rami attach to the feather sheath (Fig. 8, Stage IV). The last cells within the barb ridge to keratinize are the basilar cells adjacent to the basement membrane, forming the ventral cortex of the mature barb rami (Fig. 8, Stage IV). In this stage, a new zone of highly proliferating cells reappears along the thickened basement membrane that separates mature ridges from the pulp (Fig. 8E-F, Stage IV).

While the cell proliferation and keratinization processes in the developing barb ridges follow a similar spatiotemporal pattern to those of the rachial ridge^4,5^—occurring from the periphery toward the center along the radial axis of the pulp—our histological sections reveal two key differences between the development of barb ridges and the rachial ridge:

1. In a developing barb ridge, the first structures to keratinize are the barbule plates, whereas in developing rachial ridge, it is the dorsal cortex;
2. The dorsal cortex of the developing rachial ridge maintains constant contact with the feather sheath throughout development. In contrast, during barb ridge development, the peripheral walls (equivalent to the dorsal cortex of mature barb rami) of the developing barb rami are initially located centrally to the barbule plates and gradually move outward until they approach the feather sheath, driven by medullary expansion.

## 3. Discussion

### Taphonomy effects

Despite the identification of feathers with ventrally open rachises in several carbonized compressed fossils and Burmese amber specimens^5,26^, ventrally open barb rami have not been documented in any known avialan or non-avialan feathered dinosaurs. This raises the question of whether these unusual barb structures represent genuine morphologies or are simply preservational artifacts. Similar to the rachis, barb rami and barbules are composed of feather keratin, which is known for its resistant to water, organic solvents, and mechanical alterations including the postmortem degradation^27,28^. This durability is evident in the preservation of detailed barb and barbule structures in amber-embedded feathers^4,5,29,30^. Given this resilience, it is unlikely that barbules, being more delicate than barb rami, would be more resistant to degradation than the barb medulla and ventral cortex. The absence of the medulla and ventral cortex in these fossilized barbs, similar to the missing rachial medulla of some early feathers^5^, may be due to heterochronic developmental truncation rather than taphonomic (preservational) factors. In other words, the missing structures may result from differences in feather development in these ancient species compared to modern ones, rather than being lost through fossilization processes. This suggests that evolutionary changes in feather developmental processes, rather than preservation, could account for the absence of certain structures like the medulla and ventral cortex in fossilized feathers.

### Aerodynamic significances of ventrally open feather barbs

Unlike the ventrally open feather rachis, which has been documented not only in the fossil record but also in the summer plumage of extant penguins^5^, there is no evidence of feathers with ventrally open barbs in either fossil theropods or modern birds. This absence makes it impossible to uncover their developmental mechanisms (e.g., transcriptomic differences compared to the sandwich-like configuration of typical extant contour feathers) or to infer their potential functions. The lack of hooklets in these early feathers, as previously identified in other Burmite feathers^4,5^, combine with the presence of ventrally open feather barbs, suggest limited aerodynamic stability. Considering their actual size and structural weakness, these might be contour feathers that were covered by other covert contours or flight feathers, rendering their unstable aerodynamic performance less significant.

When airflow passes along the feather from the calamus, the instability of feathers with ventrally open barb rami (Fig. 6H) suggests that they may have resonated at certain airflow speeds. This further indicates that these early feathers were either not designed to streamline the body surface or that the animals with such feathers likely could not fly or glide. The closed vane portions of covert contour feathers in modern birds are equipped with hooklets, while the open vanes without hooklets also do not possess ventrally open barb rami^1^. Even among modern birds that have secondarily lost their ability to fly (e.g., ostriches and emus), no feathers with ventrally open barbs have been found^31^, further demonstrating that this incomplete development of barb rami has been abandoned by modern birds.

However, our functional simulation results offer insights that could inspire future biomimetic designs. The split barb rami enhance their structural stability by forming a mesh-like structure, suggesting a strategy for designing materials that need to significantly reduce weight while maintaining mechanical strength.

### Putative developmental mechanisms for bizarre feather barbs

The general developmental processes of early feathers, including cellular behaviors in early feather barb ridges, appear to be similar to those in extant feathers^4,5^. However, while altering the expressions of regulatory genes can globally affect feather structures^32^, reproducing the exact barb morphologies of early feathers in modern growing feather follicles remains challenging at present. This difficulty is compounded by the variability in barb ramus morphologies, which can differ not only within a single feather but also among various feather types. The lack of effective methods to precisely control the spatial and temporal expression of genes in developing feather follicles further complicates these efforts. In addition, the gene regulatory networks that may have produced the unique morphologies of early feathers may have decayed or been entirely lost ^4^. Consequently, the cellular and molecular events leading to the development of various ribbon-like barb rami can only be inferred from the general developmental processes observed in extant feathers, an approach traditionally employed by Paleo-Evo-Devo studies^33^.

In morphotype I feathers, barb rami can be either ribbon-like or beam-like, depending on their specific positions. Our findings suggest that the absence of the ramus medulla and ventral cortex in the ventrally open regions may result from an early truncation of cell proliferation, a failure of differentiated medullary cells to keratinize, or both, similar to previously reported ventrally open rachises. Consequently, the barb rami can only increase strength by expanding the dorsal cortex and may be the overall diameter (Figs. 5-6). If a well-keratinized medulla could be formed, the solid beam-like barb rami would not be necessary. Thus, there is no reason to assume that tissue differentiation is more advanced in the beam-like regions. We therefore propose that the solid beam consists solely of the non-expanded dorsal cortex (Fig. 9A, Bii).

**Figure 9.**
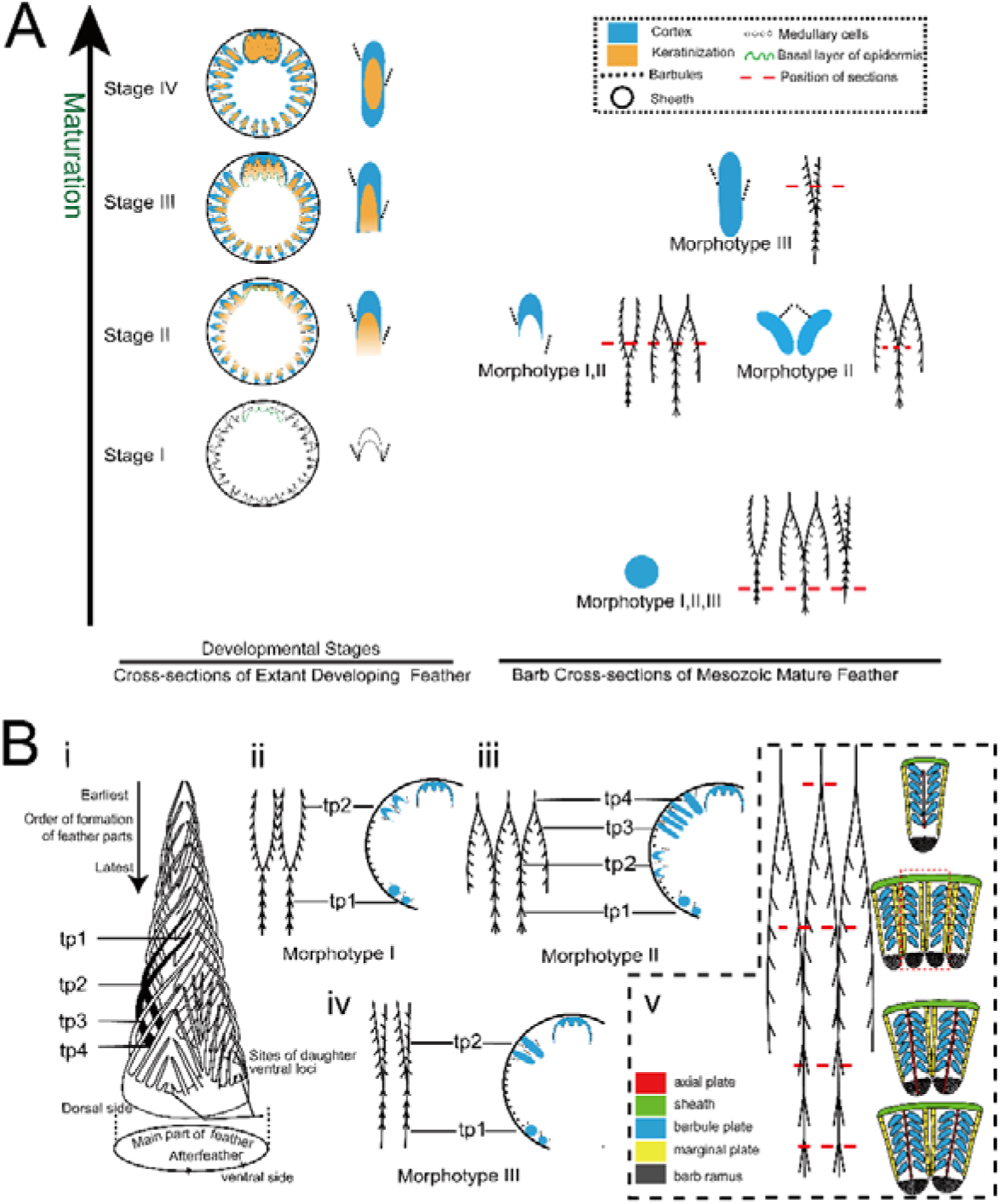
Schematic illustration showing the diverse barb morphologies resulting from incomplete developed barb rami. (A) Cross-sections of a developing extant flight feather demonstrating the morphological changes of barbs during maturation (Left), alongside cross-sections of mature barb rami in three Mesozoic feather morphotypes (I, II, and III), highlighting the diverse barb morphologies observed in Mesozoic feathers correspond to various stages of incomplete development in extant feathers (right). The red dashed lines indicate the positions where cross-sections are taken. (B) Schematic depiction of temporal developmental shifts along barb rami of each morphotype of early feather. (i) a model of growing feather with the sheath and pulp omitted, highlighting barb arrangement within feather follicle (modified from Lucas and Stettenheim^1^). The black barbs indicate those depicted in (ii-iv), illustrating that the distal tip of the barb forms first at the new barb locus while the proximal part continues to grow as they approach to the rachis (time points 1-4 are indicated as tp 1-4). (ii-iv) morphological variations along the barb rami (left) produced by temporal developmental shifts (right) for each feather morphotype. The cross-sections show only the relative positions of the barb ridges at a given time point when the corresponding part of the barb rami is formed; this does not imply that all depicted barb ridge morphologies are present in the cross-section at a given time. (v) enlarged view of barb ridges in a developing morphotype II feather, highlighting the cellular and tissue features associated with morphological changes along the barb rami.

Unlike morphotype I feathers, the morphogenic process of morphotype II feather ECNU A32 is inferred to have been more sophisticated. Like morphotype I feathers, the medulla and ventral cortex of barb rami are absent. But the split dorsal cortex suggests that the axial plate, which separates the proximal and distal barbule plates and typically terminates upon reaching the developing barb ramus, extends all the way to the apical points of the developing barb ridges throughout the morphogenic process (Fig. 9Bv). The apoptosis of the far-centrally-extended axial plates not only results in the splitting of the barb rami but also causes the basilar cells at the apexes of a single developing barb ridge to divide into two groups (Fig. 9Bv). Meanwhile, the marginal plates, located between adjacent developing barb ridges and usually extending to the apexes of the barb ridges, terminate before reaching the apical basilar cells (Fig. 9Bv). This leads to a failure in the separation of the barb rami at the apical ends of barb ridges, resulting in the fusion of halves from adjacent split rami at the barb tips (Fig. 9Bv). This outcome is further facilitated by the aggregation of the basilar cells from the adjacent developing barb ridges.

In morphotype III feathers, no trace of a medulla has been identified in the barb rami, suggesting that medullary cells did not differentiate during the morphogenesis of barb ridges. While the entire barb rami are composed solely of cortex, they do not expand transversely as seen in morphotype I feathers, but expands dorsoventrally instead (Fig. 9Bii).

Several regulatory genes involved in the normal developmental processes of extant feathers have been identified according to the well-accepted model of feather morphogenesis^4,34,35^. For example, feather regeneration relies on collar bulge region^36^, and a molecular gradient of *Wnt3a* along the rachis-new barb locus direction (equivalent to the anterior to posterior direction of Prum^3^) plays a role in the vane formation in flight feathers but not in downy feathers^37^. *Wnt* inhibitors also regulate feather regeneration and axis formation^38^. The patterning of barb ridges may be influenced by *Ephrin B1*, which is expressed in the marginal plate epithelum^39^. In a bilaterally symmetric feather, the topologies of the rachis and the barb growth zone are primarily regulated by *GDF10* and *GREM1*, with the retinoic acid signaling landscape closely related to feather asymmetry^40^. *Shh* (sonic hedgehog) and *Wnt3a* are required for the specification of marginal and axial plate cells, respectively^4,34,35^. While *Bmp4* and *Bmp2* are essential for the differentiation of barbule plates, *Shh* express is minimal in these areas^34^. *Bmp2* is transiently expressed in the peripheral marginal plates during early ramogenesis before switching to the barbule plate epithelia, indicating its role in specifying marginal plate cell fate^41^. In addition, the expression of *MMP2* in the marginal plates of feather follicle in adult chicken is involved in this process^42^. Overexpression of *Bmp4* increases barb fusion by preventing the apoptosis of marginal plate cells, whereas suppression of *Shh* could produce web-like barb fusions similar to the forked barb rami seen in morphotype II feathers^43^.

According to the well-accepted model^34^, the balance of *Noggin* and *Bmp4* determines the number, size, and spacing of barb ridges, while *Shh* is negatively regulated by *Bmps* ^34,35^. Overexpression of *Bmps* suppresses *Shh* expression and the subsequent formation of the marginal plates^34^. Conversely, suppression of *Bmps* promotes branching, likely by enhancing the ridge-forming activity of the basilar cells and specifying the fate of marginal plate cells^34^. In addition, the expression of *Ski* is also linked to the loss of the medulla in extant feathers^4^.

Based on current knowledge, ribbon-like barb rami may result from an imbalance in the expression of *Bmps* relative to their antagonists^43^. The split barb rami of morphotype II feathers could have formed through the heterotopic expression of *Shh* at the axial plate and *Wnt3a* at the marginal plates, which may have swapped the identities of these structures in certain developmental stages. From this perspective, both heterotopic (spatial) and heterochronic (temporal) expressions of these regulatory genes during feather morphogenesis could have contributed to the formation of the split, ventrally open barb rami. The absence of a medulla in morphotype III feathers might be attributed to *Ski* expression^4^. Collectively, various evo-devo mechanisms may account for the unusual barb morphologies observed in early feathers, awaiting verification as techniques become available.

### Temporal developmental shifts drive morphological variations along barb rami

Since the distal portion of a feather matures first while the proximal counterpart is still undergoing morphogenesis, any temporal physiological and/or environmental changes that trigger a phenotypic response could manifest along the proximal (immature) to distal (mature) axis of a developing feather^3,5^. This phenomenon is well demonstrated by regional morphological changes induced through the temporal expression of ectopic genes by replication-competent avian sarcoma-leukosis virus (RCAS) ^4,44^, which has been instrumental in studying feather morphogenesis. From this perspective, and based on cellular and molecular mechanisms inferred from modern feather regeneration, the temporal developmental shifts of barb rami for each feather morphotype can be described as follows:

#### Morphotype I feather barbs (**Fig. 9Bii**)

These barbs are characterized by a solid distal beam that transitions into a proximal ribbon-like, ventrally open ramus. Two developmental scenarios are proposed: (i) At the onset of barb rami formation, dorsal cortex differentiation begins, producing a solid distal beam with minimal expansion of the dorsal cortex and no differentiation of other barb ramus tissues (Fig. 9Bii, tp1); and (ii) The dorsal cortex expands in width, but medulla differentiation is absent or present but fails to keratinize, resulting in ventrally open barb rami (Fig. 9Bii, tp2). This is likely equivalent to stage II developing sandwich-like barb rami of extant feathers (Fig. 9A).

#### Morphotype II feather barbs (**Fig. 9Biii and v**)

The developmental processes of morphotype II feather barbs can be explained by three possible scenarios: (i) Basilar cells keratinized without further proliferation and differentiation, forming the solid distal beam (Fig. 9Biii, tp1); (ii) The dorsal cortex expands in width, but medulla differentiation is absent or present but fails to keratinize (Fig. 9Biii, tp2); (iii) The expanded dorsal cortex split along the midline (Fig. 9Biii, tp3); and (iv) Each branch of the split barb ramus fused to branch of the adjacent barb ramus immediately before merging into the rachis (Fig. 9Biii, tp4).

#### Morphotype III feather barbs (**Fig. 9Biv**)

These barbs are likely formed solely by the cortex without proximodistally expanded. Although no medulla is produced, this suggests that the barb rami can be fully keratinized all the way to the ventral apex. Therefore, we explain this alternative developmental pathway is more advanced compared to the solid beam and ventrally open barb rami of morphotypes I and II feathers.

Taken together, the appearance of ribbon-like barb rami offers three key evolutionary insights: **(i) Widespread absence of medulla in early feathers:** The lack of a medulla points to limited tissue differentiation within the rachis and barbs during the early stages of feather evolution. This is supported by observations that the rachis is no thicker than the feather barbs in several Burmite feathers. The inability to form well-keratinized, branching, sandwich-like structures likely meant that mechanical strengthening could only be achieved by expanding the dorsal cortex and possibly increasing the overall diameter, resulting in ventrally open rachises and barb rami. However, confirming this conclusively remains challenging due to the poorly preserved detailed structures in carbonized fossils.; **(ii) Developmental plasticity enriches feather morphology:** The presence of ventrally open regions interspersed with solid beam sections along a single barb ramus suggests that spatiotemporal developmental plasticity played a significant role in early feathers. This plasticity likely allowed for considerable variability in feather structure, contributing to the morphological diversity observed in early feathers.; and **(iii) Later evolutionary emergence of a medulla:** The consistent presence of a medulla within rachises and barb rami appears to have developed later in feather evolution, indicating that feather branching evolved before the complete stabilization of tissue differentiation within the rachis and barbs.

### Conclusions

Our findings suggests that the tissue differentiation within the rachises and barb rami of early feathers was quite limited. In some feathers that either failed to form a medulla or formed one but failed to keratinize, rachis and barb rami compensated for structural weakness by expanding the dorsal cortex. This led to the ventrally open rachises and barbs, which are rarely observed in modern feathers. The absence of a fully formed medulla and ventral cortex increased the developmental plasticity of rachises and barbs in early feathers, which likely giving rise to numerous feather morphotypes not seen in modern birds. Variations in dorsal cortex expansion and developmental plasticity of medullary tissues contributed to the morphological diversity of early feathers, in addition to the branching patterns. As the medulla and ventral cortex formed more consistently over time, the sandwich-like configuration of modern feather branches stabilized, and the developmental plasticity of the rachis and barbs gradually diminished.

## Author’s Contributions

S.W. designed the project; Y.Z., J.T. Y.W. and S.W. performed experiments; Y.Z., J.T. Y.W. C.D. and S.W. analyzed data; and S.W., Y.Z. and Y.W. wrote the paper.

## Acknowledgements

We thank Dr. Tautis Skorka and Dr. Tea Jashashvili (University of Southern California) for their assistance in scanning the specimens, and to N.D., C.C. and Y.S. (East China Normal University) for their discussion. We also appreciate the invaluable help of Y.H. (IVPP) and R.Y. (East China Normal University) in processing VG. S.W. is supported by the Human Frontier Science Program (LT000728/2018), the Zijiang Program for Talented Scholars at East China Normal University, and the Shanghai Pujiang Program (23PJ1402300).

